# A novel high-content phenotypic screen to identify inhibitors of mitochondrial DNA maintenance in trypanosomes

**DOI:** 10.1101/2020.07.16.207886

**Authors:** Migla Miskinyte, John C. Dawson, Ashraff Makda, Dahlia Doughty-Shenton, Neil O. Carragher, Achim Schnaufer

## Abstract

Kinetoplastid parasites cause diverse neglected diseases in humans and livestock, with an urgent need for new treatments. Survival of kinetoplastids depends on their uniquely structured mitochondrial genome (kDNA), the eponymous kinetoplast. Here we report development of a high-content screen for pharmacologically induced kDNA loss, based on specific staining of parasites and automated image analysis. As proof-of-concept we screened a diverse set of ∼14,000 small molecules and exemplify a validated hit as a novel kDNA-targeting compound.

## Introduction, methods, results and discussion combined

Kinetoplastids cause diverse, life-threatening diseases in humans and their livestock, namely African sleeping sickness (1), Chagas disease (2) and the leishmaniases (3) in the former and animal trypanosomiasis in the latter (4). These diseases particularly affect populations in low- and middle-income countries in many parts of the world. Currently available drugs are unsatisfactory because they cause severe, and sometimes lethal, side-effects, they are difficult to administer, and resistance continues to emerge, necessitating the development of novel anti-kinetoplastid therapies (5).

Although kinetoplastids have evolved distinct methods of infection and host immune evasion, they all share a unique biological feature: the organisation of their mitochondrial DNA (mtDNA, or kDNA in these organisms) in a peculiar structure that gave these organisms their name: the kinetoplast (6). The kDNA is intrinsically different from mammalian mtDNA, essential for parasite survival (7, 8) and a validated target for some current anti-trypanosomatid therapies (9–12), making it an attractive target for discovery of new, improved drugs (13, 14).

Drug discovery efforts are typically either phenotypic or target-based (15). While target-based campaigns have dominated efforts for decades, they often fail to produce new therapeutic molecules due to the challenge of translating promising results from reductionist biochemical and cellular assays into robust efficacy in more complex *in vivo* models (16). In contrast, phenotypic screens are often more time-consuming and expensive, and the mode(s) of action behind any identified hits are usually unknown (16). However, both approaches are complimentary and can be used synergistically to fast-track the identification of target-specific compounds that can enter the cell and reach the associated intracellular organelles to induce the desired effect. This paper describes the design, implementation and validation of a phenotypic high-content screen (HCS) with automated image analysis for the discovery of hit compounds that specifically target kDNA maintenance, using *Trypanosoma brucei brucei* (hereafter referred to as *T. brucei*) as a model system.

### HTS design and image analysis

To enable the discovery of target-specific compounds, our phenotypic screen uses a genetically engineered kDNA-independent *T. brucei* cell line which tolerates kDNA loss due to an L262P mutation in the nuclearly encoded subunit γ of the mitochondrial F1FO-ATPase (17). Non-specific cytocidal or cytostatic compounds, which would be more likely to cause side effects in the host, can readily be identified in this genetic background. Importantly, although such kDNA-independent mutants have evolved in *T. brucei* in nature (18), they have never been reported for those kinetoplastid parasites of humans and livestock that are currently responsible for by far the greatest disease and economic burden, i.e. *Leishmania* spp., *T. cruzi, T. vivax* and *T. congolense*, despite decades of use of anti-kinetoplastid compounds that affect kDNA (19).

Our HCS has been optimized for use in a high throughput 384-well format (Greiner-Bio, #781280), using a Biomek FX liquid handler (Beckman) to dilute all compounds and subsequently adding L262P *T. brucei* cells using a VIAFLO multi-well plate liquid handler (Integra) in a class II biosafety cabinet. Parasites are seeded at 50 cells per well (1 × 10^3^ cells/µL) in complete HMI-9 medium (20) supplemented with 20% (v/v) fetal calf serum and grown in an atmosphere of 5% CO2 at 37°C for 4 days (21). Following incubation, cells were stained with the cytoplasmic viability stain, 5(6)-carboxyfluorescein diacetate succinimidyl ester (CFDA-SE; CAS: 150347-59-4) at 10 μM for 15 mins at 37°C and with Hoechst 33342 nucleic acid stain at 1 μg/mL for 5 minutes at 37°C. Subsequently, cells underwent a fixation with 2% (w/v) ice-cold formaldehyde solution with vigorous mixing to avoid clumped cells, a step that is crucial for subsequent image analysis (Fig. 1A). After 24 h fixation at 4°C, cells are washed 3 times with phosphate-buffered saline by centrifuging plates at 1,000 x g for 1 min to remove any remaining dye. Cells were then transferred into 384-well plates for imaging (Greiner-Bio, #781986). The plates were centrifuged at 1,000 x g for 5 min prior to imaging acquisition at 40x magnification using an automated ImageXpress-XLS micro (Molecular Devices) HCS system. Each well was imaged across four different fields of view using DAPI (for Hoechst 33342 stain) and FITC (for CFDA-SE) filter sets. Image analysis was performed using the CellProfiler 3.1.9 software (22). Briefly, nuclear DNA and kDNA were identified based on area size of Hoechst 33342 positive objects, and viable cells were identified using the FITC channel (Fig. 1B).

**FIG 1.**
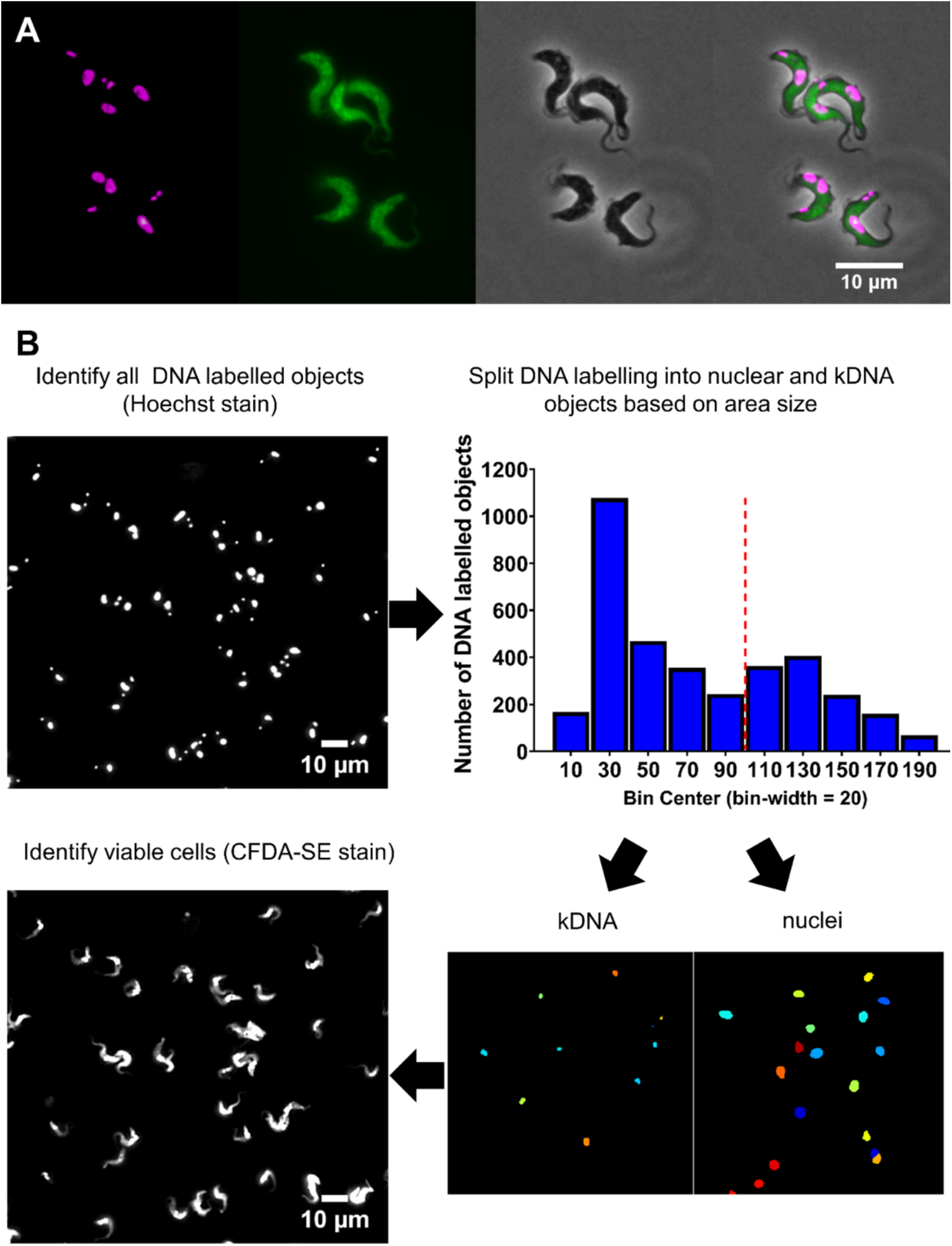
High-content screening (HCS) strategy to identify compounds inhibiting kDNA maintenance in *T. brucei*. **(A)** Representative fluorescence microscopy images of *T. brucei* using the HCS staining protocol. From left to right: Hoechst 33342 staining of trypanosome nuclei and kDNA (in magenta), CFDA-SE cytoplasmic viability stain (in green), phase contrast and merged images. **(B)** Schematic representation of the image analysis pipeline using CellProfiler. First, nuclei and kDNA were identified from the Hoechst 33342 staining (upper left panel). Next, nuclei and kDNA were identified by classifying stained objects according to area size (nuclei ≥ 100 area size, kDNA < 100 area size; bin width = 25 with bin centre ranging from 0 to 200; upper right and lower right panels). Finally, viable cells were identified using the CFDA-SE cytoplasmic viability stain (lower left panel). Each well was imaged at four different, non-overlapping positions.

### HCS performance validation and pilot screen

Plates (n=2) were prepared as above, with even-numbered columns containing a negative control (0.1% dimethyl sulfoxide (DMSO)) and odd-numbered columns containing a known inhibitor of kDNA maintenance, 10 nM ethidium bromide (EtBr) in 0.1% (v/v) DMSO, as a positive control (11). A robust *Z’* of 0.725 was calculated using the method previously described (23), indicating excellent assay performance (24).

To test the ability of our HCS to identify novel inhibitors of kDNA maintenance, 13,486 compounds were screened, from a diverse set of chemical libraries: Prestwick Chemical Library (Prestwick Chemical; 1,280 compounds), Screen-Well PKE library (Enzo Biochem; consisting of protease (53), kinase (80) and epigenetic (43) inhibitors), BioAscent 12,000 diverse chemical libraries (BioAscent Discovery Ltd.). The Prestwick Chemical library was designed to represent broad pharmacological diversity of all FDA-approved small molecule drug classes and consists of drugs with known pharmacology, toxicology and pharmacokinetic properties to support repurposing of existing drugs. The 12,000 BioAscent compound library is a subset representing the chemical diversity of a 125,000 parent compound library, enabling subsequent expansion of screening hits to explore structure-activity relationships. All compounds were screened at a final concentration of 10 μM in 0.1% (v/v) DMSO in a 384-well format, where the first four columns had alternating positive (EtBr) and negative (DMSO) controls (Fig. S1). Additionally, the PKE and Prestwick Chemical libraries were screened at a final concentration of 1 μM. The screens were performed in 5 batches (48 plates in total), with an robust *Z’* (25) ranging from 0.63 to 0.9 between batches. The HCS identified 152 compounds with a reduced ratio of kDNA per nucleus (*Z*-score < -2; Fig. 2A).

**FIG 2.**
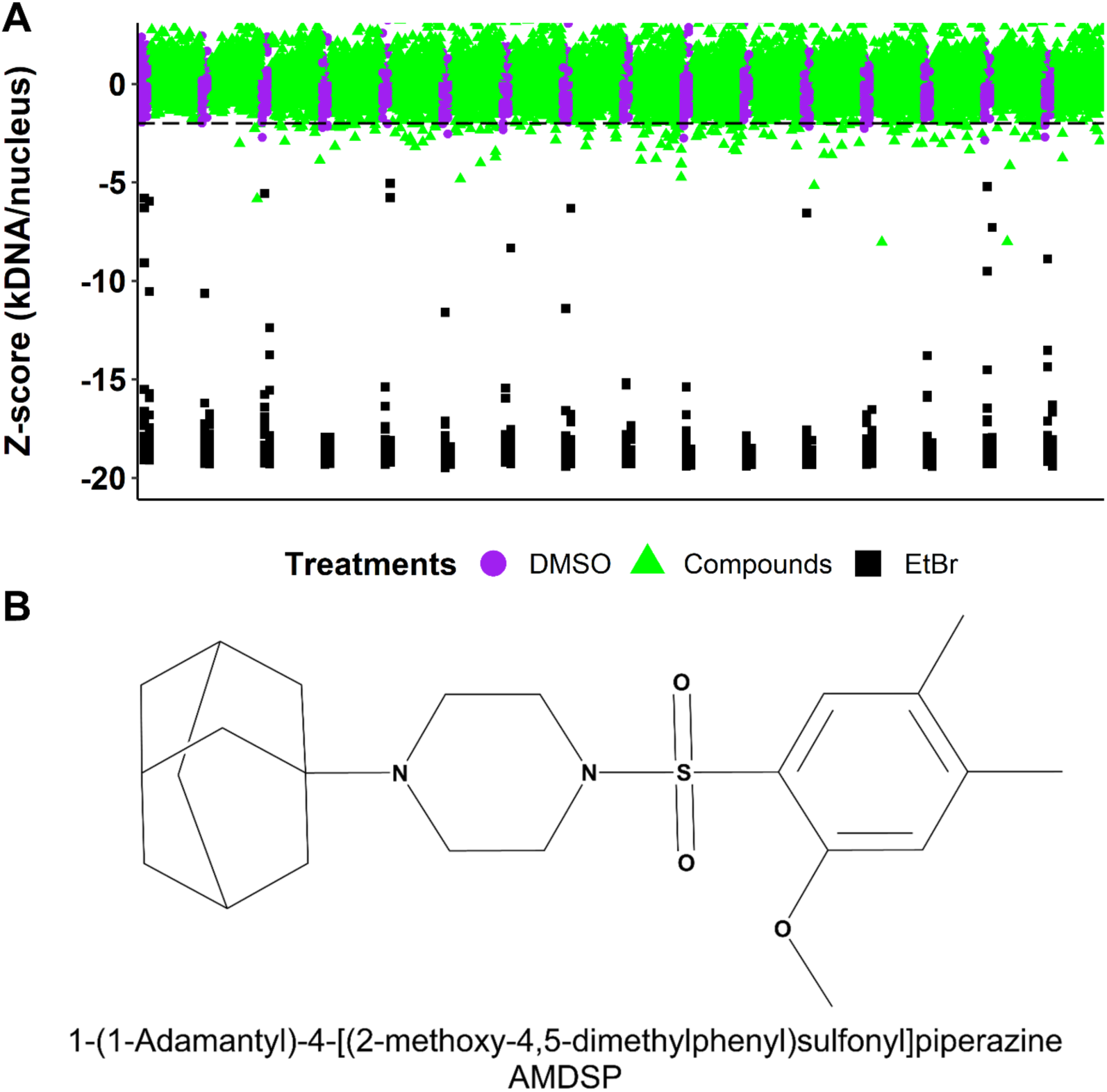
HCS hit selection. **(A)** Tested compounds were ranked based on the decrease of kDNA/nucleus ratio in imaged wells (*Z*-score < -2 (dashed black line)). Images of the top 50 hits were then re-examined using ImageJ software and the potency of 10 compounds with biggest decrease in kDNA/nucleus ratio was tested in WT (kDNA-dependant) and L262P (kDNA-independent) *T. brucei* cells. **(B)** Structure of the validated kDNA targeting compound AMDSP (BCC0052412).

### Hit validation

For the top 50 compounds, based on ranking by *Z*-score, we reviewed the microscopy images and manually recounted kinetoplasts and nuclei using ImageJ software (Table S1). The top 10 candidates were chosen based on decrease of kDNA to nucleus ratio and subsequently purchased from commercial suppliers. Compounds were dissolved in DMSO, and their potency against kDNA-dependent (‘WT’) and -independent (‘L262P’), but otherwise isogenic, *T. brucei* cells was evaluated using an adapted 3-day Alamar Blue method (17). Specific inhibitors of kDNA maintenance will preferentially kill kDNA-dependent *T. brucei*, with the most specific compound reported to date being EtBr, with a selectivity index of ∼300 (11). One compound, 1-(1-Adamantyl)-4-[(2-methoxy-4,5-dimethylphenyl)sulfonyl]piperazine (AMDSP, BioAscent code BCC0052412) reproducibly affected the viability of WT *T. brucei* cells at lower concentration compared to L262P cells (Fig. 3A). The IC50 for WT cells was 1.9 μM, while the IC50 for L262P cells was estimated to be in the range of 8 μM (the value could not be determined more precisely due to poor compound solubility in DMSO at higher than 12.5 mM stock concentration). We also observed a significant reduction of kDNA area in AMDSP-treated cells compared to control cells (Fig. 3B). To investigate the time required for AMDSP to induce death, we performed growth curves in WT and L262P cells at a final compound concentration of 12.5 μM in 0.1% (v/v) DMSO (Fig. 3C and 3D). AMDSP treatment resulted in growth inhibition of WT cells after 3 days and cell death after 4 days, while L262P cells survived. These data are consistent with the dynamics of other trypanocidal compounds that preferentially target kDNA (e.g. EtBr). Altogether, these data confirm that an important part of the mode of action of AMDSP in trypanosomes is interference with kDNA maintenance. Similarity searches of the full BioAscent library suggest up to 150 related compounds that could be tested against *T. brucei* in the future to explore structure-activity relationships.

**FIG 3.**
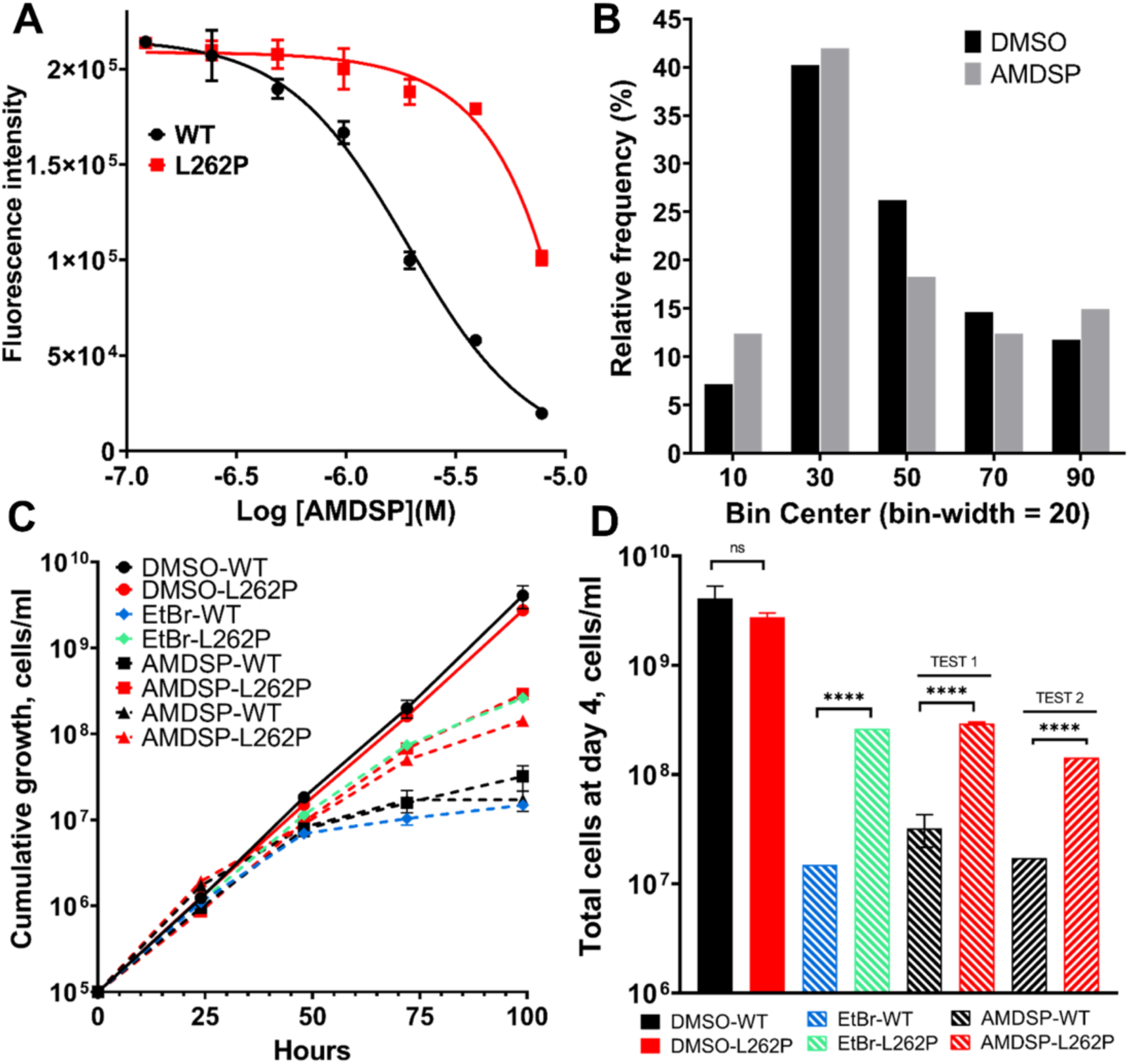
Hit validation. **(A)** Dose-response curves of AMDSP induced trypanosome death for kDNA-dependant (WT, black squares) and kDNA-independent (L262P, red squares) cells. **(B)** Relative frequency distribution of kDNA area calculated using the CellProfiler pipeline in trypanosomes treated with AMDSP (grey, n=274) and DMSO (black, n=656). Bin width = 20. The frequency distribution statistical significance was assessed using Kolmogorov-Smirnov test; *p* = 0.0076. **(C)** Cumulative growth curves of *T. brucei* cells cultured in the presence (dashed lines) and absence (solid lines, filled circles) of 12.5 µM AMDSP or 10 nM EtBr. **(D)** Comparison of cumulative cell numbers in **(C)** after 96 h between WT and L262P cells. *t*-test, *p* < 0.00005 (****).

AMDSP is composed of piperazine, benzene and adamantane rings with a tertiary sulfonamide group. Adamantane derivatives, such as the well-studied drug, amantadine (1-amino-adamantane), show good pharmacokinetics in humans and have been used as anti-viral and anti-parkinsonian agents (26). Moreover, the discovery of amino-adamantane derivatives with enhanced trypanocidal activity (27) has spurred efforts for the development of more potent adamantane-benzene derived trypanocidal agents (28). Piperazine-based anti-helminthic drugs (29) have also gained interest in drug design studies because of their trypanocidal activity (30). The exact mechanism(s) by which the described derivatives affect kinetoplastids remains unknown but, based on our findings, effects on kDNA should be explored.

### Identification of other anti-trypanosomatid compounds with unknown mode of action

In addition to a novel inhibitor of kDNA maintenance, we also identified compounds that strongly affected the viability of the kDNA-independent *T. brucei* cell line used for screening and that therefore must act via a different mechanism. To find such trypanocidal or trypanostatic hits, we first corrected for positional growth effects in our plates using the median polish normalisation method (31) (Fig. S1). We then scored for hits affecting *T. brucei* viability based on less than 10 total nuclei per image with *Z*-scores < -2 or < -1. We identified 279 and 337 hits, respectively, corresponding to hit rates of 2.1% and 2.5% (Table S2; Figure S2). These include 31 compounds (*Z*-score < -2) from the Prestwick Chemical Library that inhibit trypanosome growth at 10 μM and 1 μM (underlined in Table S2), suggesting a good starting potency for any lead development efforts. Incidentally, among the compounds tested in our proof-of-concept screen were 9 compounds with known anti-trypanosomatid activity (32). Seven of these compounds were among the hits with a *Z*-score < -1 (Fig. S2). This further confirms the robustness of our HCS assay and suggests that this assay could also be used as a general phenotypic screen for the identification of compounds with anti-trypanosomatid activity. Robustness of the screen for the latter purpose could be improved further by addressing the considerable plate edge effect that we observed (Fig. S1).

In conclusion, we successfully established and validated a kDNA maintenance based phenotypic HCS with automated image analysis, using an engineered kDNA-independent *T. brucei* cell line as a kinetoplastid model system. A proof-of-concept screen of diverse small compound libraries identified and validated a novel compound affecting kDNA maintenance in *T. brucei*. Furthermore, we identified other anti-trypanosomatid compounds with activity in the low micromolar range (but with unknown molecular targets) that could be useful starting points for trypanosomatid drug development.

## Acknowledgements

M.M., J.D., D.D-S., N.O.C., A.M., and A.S. designed the research; M.M., J.D., A.S. analysed the data; M.M. performed the research; M.M. and A.S. wrote the paper.

We thank Zandile Nare for helpful discussions and Angus Morrison (BioAscent) for suggestions on BCC0052412 analogs.

This work was supported by Senior Non-Clinical Fellowship MR/L019701/1 from the UK Medical Research Council to A.S and Institutional Strategic Support Fund (ISSF3) award (reference IS3-R2.28) to A.S. for salary to M.M. and consumables.

